# Functional Molecular Switches of Mammalian G Protein-Coupled Bitter-Taste Receptors

**DOI:** 10.1101/2020.10.23.348706

**Authors:** Jérémie Topin, Cédric Bouysset, Jody Pacalon, Yiseul Kim, MeeRa Rhyu, Sébastien Fiorucci, Jérôme Golebiowski

## Abstract

Bitter taste receptors (TAS2Rs) are a poorly understood subgroup of G protein-coupled receptors (GPCRs). The experimental structure of these receptors has yet to be determined, and key-residues controlling their function remain mostly unknown. We designed an integrative approach to improve comparative modeling of TAS2Rs. Using current knowledge on class A GPCRs and existing experimental data in the literature as constraints, we pinpointed conserved motifs to entirely re-align the amino-acid sequences of TAS2Rs. We constructed accurate homology models of human TAS2Rs. As a test case, we examined the accuracy of the TAS2R16 model with site-directed mutagenesis and *in vitro* functional assays. This combination of *in silico* and *in vitro* results clarify sequence-function relationships and identify the functional molecular switches that encode agonist sensing and downstream signaling mechanisms within mammalian TAS2Rs sequences.

**Classification:** Biological sciences, Computational biology, and bioinformatics

## Introduction

Bitterness is one of the basic taste modalities detected by the gustatory system. It is generally considered to be a warning against the intake of noxious compounds^1^ and, as such, is often associated with disgust and food avoidance^2^. At the molecular level, this perception is initiated by the activation of bitter taste receptors. In humans, 25 genes functionally express these so-called type 2 taste receptors (TAS2Rs), which provide the capacity to detect a wide array of bitter chemicals^3^. Further, TAS2Rs are also ectopically expressed in non-chemosensory tissues, making them important emerging pharmacological targets^4–6^.

TAS2Rs are G protein-coupled receptors^7^ (GPCRs) classified as distantly related to class A GPCRs. They were previously classified with class F GPCRs^8^ and more recently as a separate sixth class evolved from class A^9,10^. The sequence similarity between TAS2Rs and class A GPCRs is in the range of 14%-29%^11^. Structure-based sequence alignment has placed TAS2Rs in the class A family, which contains the olfactory chemosensory receptors sub-family^12^. TAS2Rs have been recently labelled as class T in the GPCR database (GPCRdb) (Fig. 1a)^13^.

**Figure 1.**
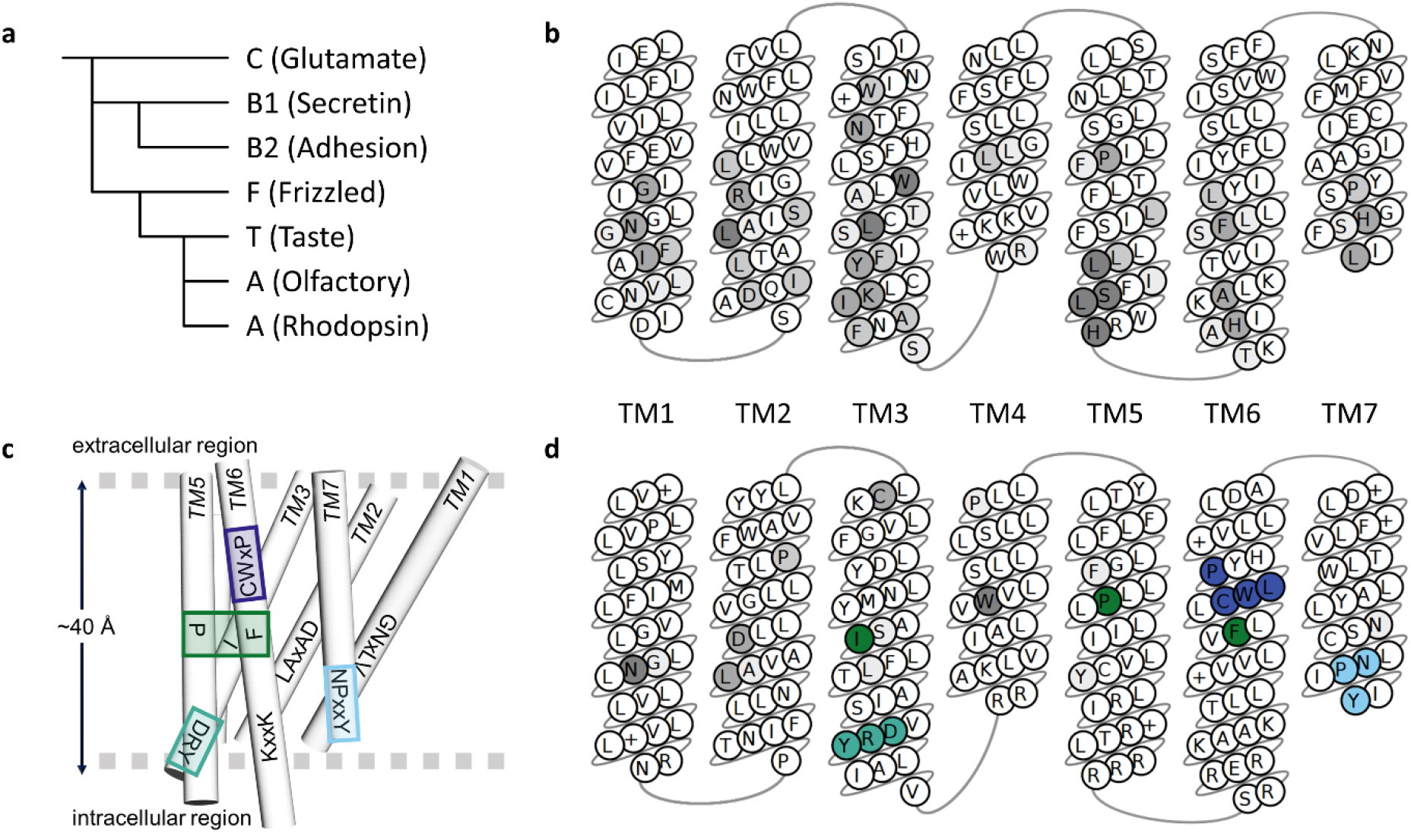
a) Schematic phylogenetic tree of GPCR classes according to Cvicek et al.^12^. b) Snake plot representation of transmembrane segments (TM) of mammalian TAS2Rs consensus sequences, colored in grey scale according to sequence conservation. c) Non-olfactory class A GPCR sequence hallmarks (transmission switch in blue, hydrophobic connector in green, ionic lock in sea green, hydrophobic barrier in light blue). d) Snake plot representation of non-olfactory class A GPCR consensus sequences.

Structurally, GPCRs are made up of seven transmembrane (TM) helices named TM1 to TM7 that form a bundle across the cell membrane. How GPCRs achieve specific robust signaling and how these functions are encoded in their sequences are pending fundamental questions. GPCR activation relies on so-called molecular switches, which allosterically connect the ligand binding pocket to the intracellular G protein coupling site in order to trigger downstream signaling^14^. In class A GPCRs (including olfactory receptors, ORs), these molecular switches consist of conserved sequence motifs (Fig. 1c). The “toggle/transmission switch” CWxP^TM6^ (or FYGx^TM6^ in ORs) senses agonist binding. The other motifs, which propagate the signal, include the “hydrophobic connector” PIF^TM3-5-6^, the NPxxY^TM7^, the “ionic lock” DRY^TM3^, and a hydrophobic barrier between the last two^15–18^.

To date, experimental structures have not been determined for any TAS2Rs, but the following hallmark motifs have been defined based on sequence conservation: NGFI^TM1^, LAxSR^TM2^, KIANFS^TM3^, LLG^TM4^, PF^TM5^, HxKALKT^TM6^, YFL^TM6^, and PxxHSFIL^TM7^ ^7^. These conserved motifs are highly dissimilar between TAS2Rs and class A GPCRs (Fig. 1b,d and Table 1), leading to different sequence alignments. The main discrepancies occur in TM3, TM4, TM6, and TM7^11,19–30^, making it difficult to infer TAS2R functional molecular switches. These discrepancies remain a central issue in understanding the complex allosteric TAS2R machinery. The present study aims to identify the molecular switches that control TAS2R functions. We present an integrative protocol that advances comparative modeling of TAS2Rs. Case studies of site-directed mutagenesis followed by *in vitro* functional assays on human TAS2R16 then evaluated the roles of the predicted molecular switches in TAS2Rs.

**Table 1.** Key residues and consensus motifs. Superscripts refer to the Ballesteros–Weinstein numbering scheme.

| TM | class A GPCR | OR | TAS2R |
| --- | --- | --- | --- |
| 1 | GN<sup>1.50</sup>xLV | GN<sup>1.50</sup>LLI | N<sup>1.50</sup>GFI |
| 2 | LAXAD<sup>2.50</sup> | LSFXD<sup>2.50</sup> | LAXSR<sup>2.50</sup> |
| 3 | L<sup>3.43</sup> | L<sup>3.43</sup> | L<sup>3.43</sup> |
|  | DR<sup>3.50</sup>Y | MAYDR<sup>3.50</sup>YVAIC | K<sup>3.50</sup>IANFS |
| 4 | W<sup>4.50</sup> | W<sup>4.50</sup> | L<sup>4.50</sup>LG |
| 5 | P<sup>5.50</sup> | P<sup>5.50</sup>F | P<sup>5.50</sup>F |
|  | Y<sup>5.58</sup> | Y<sup>5.58</sup> | F<sup>5.58</sup> |
| 6 | K<sup>6.32</sup>xxK | RxK<sup>6.32</sup>AFSTC | HxK<sup>6.32</sup>ALKT |
|  | CW<sup>6.48</sup>LP | FY<sup>6.48</sup>G | YF<sup>6.48</sup>L |
| 7 | SxxNP<sup>7.50</sup>xxY | PxxNP<sup>7.50</sup>xIY | PxxHS<sup>7.50</sup>FIL |

## Methods

### Sequence alignment

Automatic multiple sequence alignment (MSA) of TAS2Rs was performed with class A and class F templates (labelled *ClustalO* and *classF,* respectively) using ClustalO^31^ with default settings in the Jalview interface (v2.11.0)^32^. These MSAs were not modified. Another MSA, labelled *Chemosim,* was completed using class A templates, 339 class II ORs and TAS2Rs. The *Chemosim* alignment was then manually refined using constraints from functional assays in the literature (as described in the results section). We specifically focused on the 339 class II ORs because they contain relevant motifs for TAS2Rs alignment and because TM sequence conservation is higher than in a mixture of class I and class II human ORs. TM segments were predicted by the PPM webserver^33^. The final *Chemosim* MSA is provided as an additional supporting file (TAS2R-OR-templates.pir).

### Template selection for comparative modeling of bitter taste receptors

Class A GPCR templates were selected by submitting each of the 25 human TAS2Rs UniprotKB accession numbers to the Swiss-Model modeling server^34^. From the proposed templates for human TAS2Rs, 46 with at least 10% sequence identity were kept. Templates were then grouped by protein name and sorted by resolution and average sequence identity with TAS2Rs. The highest resolution template from each group was retained, resulting in 19 templates. Finally, six GPCR class A templates were selected to maximize structural diversity. As TAS2Rs have been suggested to be part of the same family as the frizzled receptors^35^, 3 class F GPCR templates were also considered: the human FZD4 receptor^36^ and 2 structures of the human SMO receptor^37^. The PDB code for the six class A templates were as follows: rhodopsin (6FUF)^38^, β1-adrenergic (4BVN)^39^, β2-adrenergic receptor (5JQH)^40^, angiotensin II type 1 (4YAY)^41^, chemokine receptor CXCR4 (3ODU)^42^, serotonin receptor 5-HT2C (6BQG)^43^.

### Integrative structural modeling of TAS2R

Using the protocols described above (*Chemosim*, *Gomodo*, *ClustalO*, *GPCRdb*, *BitterDB*, and *classF*), we built a large number of 3D models and evaluated and ranked them using a meta-score defined as the average of the pocket and helicity score (Fig. 2). This score provides a unique descriptor that accounts for both GPCR structural requirements and TAS2R experimental constraints.

**Figure 2.**
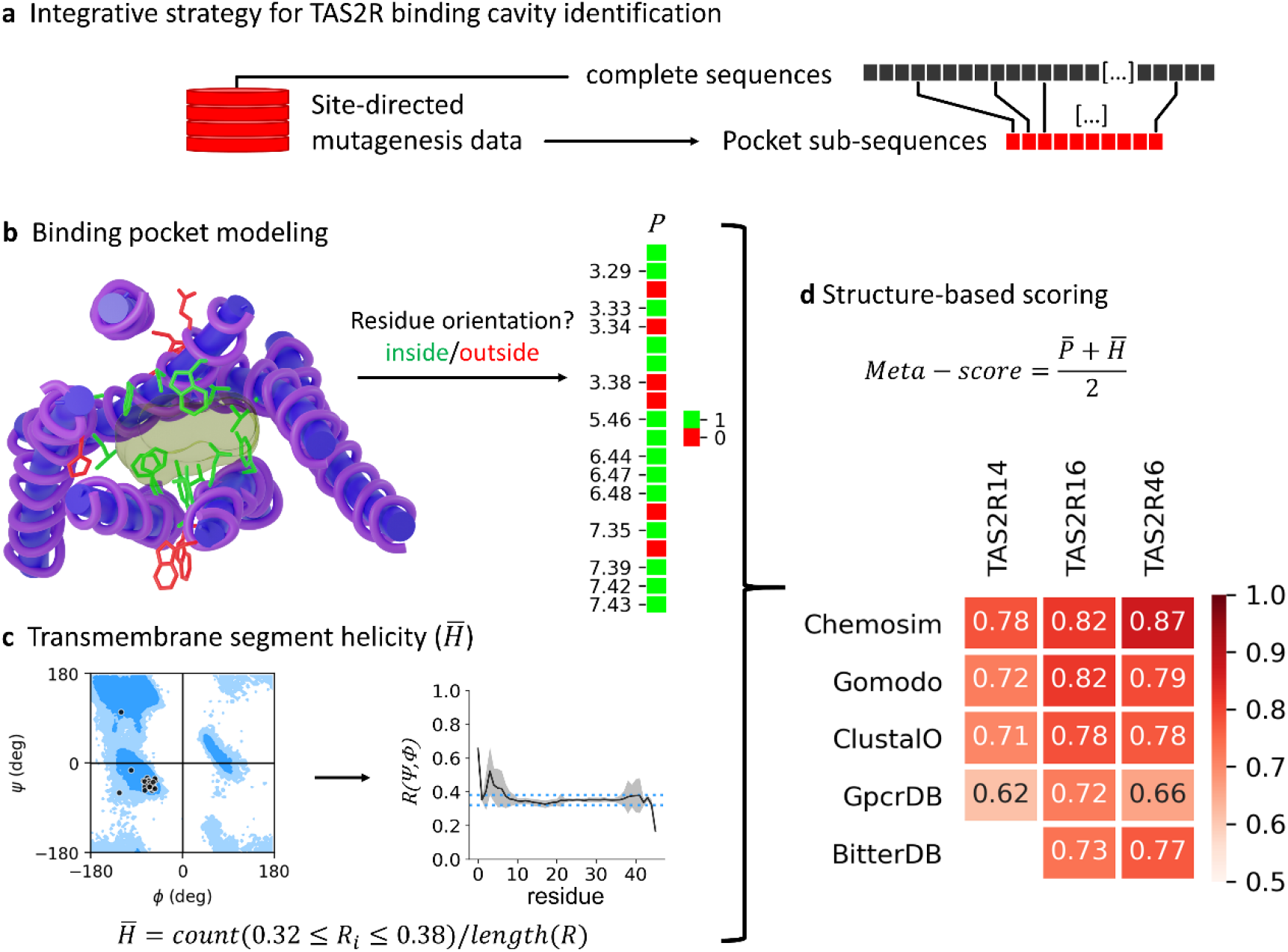
a) An integrative approach to identify the TAS2R binding pocket that is used as a constraint in comparative modeling with the Chemosim protocol. b) A pocket fingerprint was extracted based on the positions of binding residues in the 3D model. The light brown surface represents the binding pocket. c) The helicity of the TM segment was analyzed and d) combined with the pocket fingerprint to calculate a structure-based normalized meta-score. The meta-scores of the best 3D models of TAS2R14, 16 and 46 structures generated by the different comparative modeling protocols are shown in panel d).

For each alignment (*ClustalO*, *Chemosim,* and *classF*) and each template, we generated 1000 homology models using Modeller v9.21^44^ with a maximum of 300 conjugate gradient minimization steps and refinement by molecular dynamics with simulated annealing (“md_level”=slow). The remaining parameters were set to default from the “automodel” class. The BitterDB and GPCRdb webservers provided additional 3D models of each TAS2R. The GOMoDo^45^ webserver was also used to automatically generate models of TAS2Rs based only on the sequence (labelled *Gomodo* in the analysis). Default options were used, excepting the number of models which was set to the maximum (99 models).

#### Evaluation of the model pocket score

To identify residues oriented toward the binding pocket, the following protocol was implemented in Python: i/ For each of the 25 human TAS2Rs, a reference 3D model was selected from the *Chemosim* models. All reference models were then structurally aligned to the TAS2R16 reference. ii/ A unique grid of points broadly covering the binding site of class A GPCRs was generated and aligned to the coordinates of the TAS2R16 reference. iii/ Each TAS2R model was aligned to its reference based on the alpha carbons of the TM residues. iv/ Residues whose sidechain center of mass (SCM) was within 8.0 angstroms of any grid point, and whose angle between the SCM, the alpha carbon, and any grid point was lower or equal to 30 degrees, were considered as oriented towards the pocket. Only residues annotated as involved in ligand binding were kept (see supporting file TAS2R-msa_annotated.xlsx). v/ The pocket score was calculated as the fraction of residues oriented towards the pocket for each TM, averaged across all TMs. 3D structure alignment was performed with MDAnalysis v1.0.0^46^, and distance and angle calculations were performed with scipy v1.5.0^47^ and numpy v1.19.0^48^.

#### Evaluation of TM helicity

The Ramachandran number49 (*R*) was used to check the structural quality of the TM domains of each model produced. *R*, which is based on the ϕ and Ψ dihedral angles, can be seen as a short numerical form of the Ramachandran plot. First, we analyzed the helicity of 358 class A GPCR X-ray structures to set the experimental range and found an average value of 0.35. Thus, a residue was considered in an alpha-helix conformation if its *R* value fell between 0.32 and 0.38.

To discard misshapen 3D models having severe kinks in the middle of TM domains, we introduced a function based on *R*. We defined the function *f*(*r*) = count(|*r*_*i*_ − *R*_*ref*_| ≤ *σ*), where *r* is a moving subset of six consecutive *R* values that are shifted forward until all *R* values for a given TM helix have been sampled; *R*_*ref*_ = 0.35 is the average *R* value based on X-ray structures; and *σ*=0.07 is a parameter that was optimized to exclude misfolded TM proteins while keeping X-ray structures. If at any point the result of *f*(*r*) was lower than 4 for any TM residue, the model was discarded. A helicity score 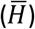 was then calculated as the fraction of TM residues satisfying the condition: 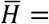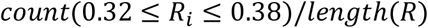. Among all considered X-ray structures, the minimum 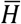 value obtained was 0.789. This threshold was used to filter out irrelevant models.

#### Assessing meta-score accuracy

The meta-score was defined as the average of the pocket and helicity scores. The relevance of the meta-score was assessed by building a homology model of the human smoothened receptor (class F) from a β2-adrenoceptor template (class A, with a low shared sequence identity [9%] with class F, PDB 5JQH). Using the experimental structure of a human smoothened receptor (PDB 4JKV), the RMSD of the best model was then calculated from the meta-score or from the scores available in Modeller or the QMEANBrane^50^ webserver. As shown in Fig. S2, the meta-score outperformed classical metrics when ranking GPCR models based on distantly related GPCR templates.

### Cell culture and transfection

Plasmids encoding TAS2R16 and G16αgust44 were constructed as previously described 51. G16_αgust44_ and TAS2R16 were cloned into a CMV promoter-based vector and expressed constitutively. Point mutations on the TAS2R16 clone were obtained from a commercial service (Macrogen Inc., Seoul, Republic of Korea), which also performed DNA sequencings of the mutant genes. The TAS2R16 and G16_αgust44_ expression plasmids were co-transfected (4:1) into HEK293T cells using Lipofectamine 2000 (Invitrogen, Carlsbad, CA, USA). Cellular responses were measured 18–24 h after transfection. Cells were cultured at 37°C in a humidified atmosphere of 5% CO_2_. The culture medium was Dulbecco’s modified Eagle’s medium (DMEM) supplemented with 10% heat-inactivated fetal bovine serum (FBS), 100 IU/ml penicillin G, 100 μg/ml streptomycin, 2 mM L-glutamine, and 1 mM sodium pyruvate (Invitrogen).

### Quantitative measurement of intracellular Ca^2+^ in bitter taste receptors upon stimulation with salicin

The compound-induced changes in cytosolic Ca^2+^ concentrations were measured using a FlexStation III microplate reader (Molecular Devices, Sunnyvale, CA, USA). Cells transfected with TAS2R16 were seeded onto 96-well black-wall CellBind surface plates (Corning, NY, USA). After 18–24 h seeding, the cells were washed with assay buffer (130 mM NaCl, 10 mM glucose, 5 mM KCl, 2 mM CaCl_2_, 1.2 mM MgCl_2_, and 100 mM HEPES; pH 7.4) and incubated in the dark, first at 37°C for 30 min, and then at 27°C for 15 min in assay buffer consisting of Calcium-4 (FLIPR Calcium 4 Assay Kit, Molecular Devices). After the samples were treated, the cell fluorescence intensity (excitation, 486 nm; emission, 525 nm) was measured. The results were plotted with ΔF/F^0^ on the y-axis, where ΔF is the change in Calcium-4 fluorescence intensity at each time point, and F^0^ is the initial fluorescence intensity. The responses from at least three wells (*n* = 3) with the same stimulus were averaged.

## Results and discussion

### Matching conserved motifs between Class A GPCRs and TAS2Rs

The prediction of TAS2Rs tertiary structure based on sequence similarity remains challenging due to discrepancy in the published alignment^11,19–30^. We have already shown that refining the sequence alignment of ORs with non-olfactory class A GPCRs by including site-directed mutagenesis produces relevant three-dimensional models of chemosensory receptors. These models have been supported by a large amount of experimental data16,18,52,53. We thus apply a similar integrative strategy to TAS2Rs. To overcome the lack of sequence similarity between TAS2Rs and GPCRs with known structures, we inserted 339 human class II OR sequences in the alignment. Subsequent manual data curation involved integration of site-directed mutagenesis data from the literature for 136 amino-acids positions, i.e. 45% of the entire TAS2Rs sequence (see ESI TAS2R-msa_annotated.xlsx). Our alignment (Fig. S1) highlights the key residues and consensus motifs in all human TAS2Rs, which correspond to the functional molecular switches in ORs and non-olfactory class A GPCRs (Fig. 1b,d). They are detailed above and summarized in Table 1.

TM1, 2 and 4 did not contain motifs involved in downstream signaling. In TM1, the NGFI^TM1-TAS2R^ motif corresponds to GNLLI^TM1-OR^ in OR and GNxLV^TM1-classA^ in non-olfactory GPCR templates (see Fig. S1). In TM2, R^2.50-TAS2R^ in the LAxSR^TM2-TAS2R^ motif aligns with D^2.50-OR/classA^, which in class A GPCRs constitutes a sodium ion binding site that stabilizes inactive receptor conformations^54^. Position 2.50 in TAS2Rs is positively-charged and unlikely to be involved in sodium binding. Rather, it is hypothesized to stabilize the structure of TAS2Rs.^21^ The sequence alignment of TM4 was not straightforward, as it lacks the canonical W^4.50-OR/classA^. The highly conserved leucine L^4.50^ of the LLG^TM4-TAS2R^ motif aligns with the most conserved W4.50-OR/class A.

TM3, 5, 6, and 7 contained functional molecular switches which have been identified in class A GPCR experimental structures^14^.

In TM3, K^3.50^ in the KIANFS^TM3-TAS2R^ motif matches R^3.50^ of the DRY^TM3-classA^ and MAYDRYVAIC^TM3-OR^ motifs. The DRY motif constitutes the ionic lock in ORs and non-olfactory class A GPCRs. This also aligns the highly conserved L^3.43^, with a leucine found at position 3.43 in both non olfactory class A GPCRs and OR (Table 1).

In TM5, the conserved P^5.50^ of the PF^TM5-TAS2R^ motif corresponds to the PF^TM5-OR^ and P^TM5-classA^ motifs/residue involved in the so-called “hydrophobic connector” (P^5.50^I^3.40^F^6.44^ in class A GPCRs). Another conserved aromatic residue that is found in 52% of TAS2Rs, F^5.58^, consistently aligns with the conserved Y^5.58^ known to be important for GPCR activation^18,55^.

In TM6, the HxKALKT^TM6-TAS2R^ motif matches both a comparable motif in non-olfactory class A GPCRs and the typical OR motif RxKAFST^TM6-OR^. The “toggle/transmission switch” (CW^6.48^LP^classA^ and FY^6.48^G^OR^) aligns with the YF^6.48^L motif in TAS2Rs. The location of this YF^6.48^L motif at the bottom of the pocket is consistent with site-directed mutagenesis results, suggesting a ligand-sensing role, as is the case for class A GPCRs^16,56^.

The extracellular part of TM7 is well-documented to belong to the ligand binding pocket in TAS2Rs and other GPCRs^20,24,56^. This is consistent with its high sequence variability (see Fig. S1). TM7 intracellular residues show higher conservation, as they are involved in GPCR signaling^16,56^. These conserved motifs, however, show little similarity between TAS2Rs and other GPCRs. Here, the comparison with ORs is highly instructive: from the P^7.46^xLNP^7.50^xIY^TM7-OR^ motif found in ORs, P^7.46^ is shared with TAS2Rs, and NP^7.50^xxY is found in other class A GPCRs. P^7.46^ and P^7.50^ are conserved in 76% and 28% of human TAS2Rs, respectively. The PxxHSFIL^TM7-TAS2R^ motif is consequently aligned with PxLNPxIY^TM7-OR^, which itself matches the highly conserved xxxNPxxY^TM7-classA^ motif^20^.

### Predicted tertiary structure of TAS2Rs

Based on this refined alignment, we tested various protocols and structural templates to build accurate 3D homology models of TAS2Rs. Among the TAS2Rs, receptors TAS2R14, 16, and 46 were selected to evaluate the approach, as previous work on these receptors involving site-directed mutagenesis provides data to determine the residues within their binding pocket. According to our meta-score, the best models of these three receptors were obtained using the *Chemosim* approach and a single template, either the β2-adrenoceptor (PDB 5JQH) or the β1-adrenoceptor (PDB 4BVN) structure (Fig. 2 & S3). The performance of each protocol is compared in Fig. S3 and S4. *Gomodo* and *ClustalO* approaches led to comparable models, with slight improvement over *BitterDB* and, in most cases, substantial improvement over *GPCRdb*. The use of class F templates systematically led to models with misfolded helices (Fig. S4).

These models and analysis were then extrapolated to the full human TAS2Rs repertoire. Even if limited experimental data is available, we were able to define a consensus TAS2R cavity based on the positions identified simultaneously in TAS2R14, 16 and 46. We also extended the definition of a specific TAS2R cavity to residues identified by site-directed mutagenesis. The best models for the entire TAS2R family were obtained using GPCR templates in their closed conformation (Fig. S6), with the exception of TAS2R38, for which the open-conformation 5-HT2C receptor (PDB 6BQG) was best. On average, the templates 5JQH, 4BVN, and 2RH1, all of which correspond to adrenergic receptors, performed best. In this study, we found no relationship between the performance of the protocols and the percentage sequence identity of the templates used to build the models. At 10–15%, the sequence identity between TAS2Rs and class A templates is too low to be a discriminating criterion.

The best *Chemosim* model obtained for each human TAS2R is provided as a PDB file in the supporting information. Projecting TAS2Rs sequence conservation onto the 3D structure showed that the models retain the structural characteristics of the GPCR (Fig. S5). The most conserved residues were located in the intracellular region of the receptor that binds the G protein, while the greatest variability was found in the extracellular ligand-binding pocket. Analysis of the binding cavity (Fig. S7) revealed high diversity within the hTAS2Rs family. The pocket volume ranged up to 400 Å^3^ and 700 Å^3^ for hTAS2R13 and hTAS2R39, respectively, corresponding to the structural features of a GPCR^57^. Although no obvious structure-function relationship was revealed by the analysis of the cavity volume, the hydrophobicity partially correlated with the receptor range of response. The binding cavities of TAS2Rs with broad ligand spectrums tended to be more hydrophobic than those of narrow-spectrum receptors (Fig. S7), consistent with previous studies showing a correlation between hydrophobicity and GPCR promiscuity^58,59^.

### Evaluating the function role of molecular switches

To evaluate the functional role of the predicted molecular switches, twelve residue positions on TAS2R16 were subjected to site-directed mutagenesis followed by *in vitro* functional assays with salicin (Fig. 3 and Table S2). The residues mostly belonged to TM3 and TM6, which, in GPCRs, are well-known to be involved in agonist sensing and activation14.

**Figure 3.**
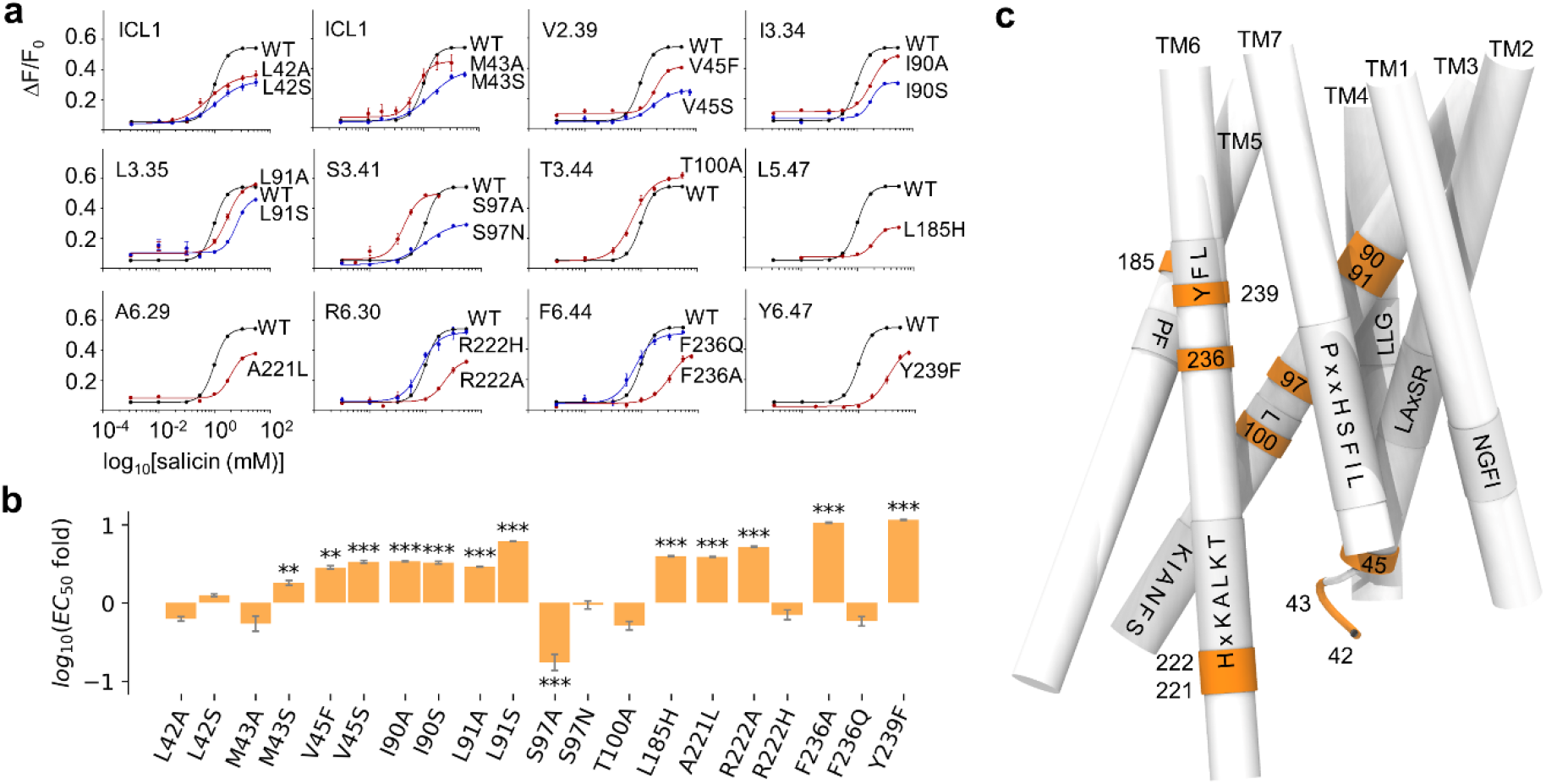
a) *In vitro* functional assays of wild-type (WT) TAS2R16 and single-point mutants stimulated by salicin. b) EC50 fold (compared to WT) expressed as log(EC50(MUT)/EC50(WT)) for the twenty TAS2R16 mutants considered in this study. Positive values indicate a reduced response to salicin in the mutated receptor compared to the WT. *** p < 0.001, ** p <0.01, and * p < 0.05 versus the WT group (one-way ANOVA followed by Dunnett’s test). c) Representative structure of TAS2R16 highlighting the location of the mutated residues. The TM domains are presented as sticks. The positions of mutated residues are colored in orange, and the molecular switches revealed by the sequence alignment are indicated on the structure.

Using our model as a basis, we investigated residues found in the ligand binding pocket (90^3.35^, 91^3.36^, and 185^5.47^) and at or around the predicted molecular switches (45^2.39^, 97^3.41^, 221^6.29^, 222^6.30^, 236^6.44^, and 239^6.47^). Residues 42^ICL1^, 43^ICL1^, and 100^3.44^ were predicted to be far from the molecular switches. All mutants showed a specific, dose-dependent response to salicin (Fig. 3), confirming that they are expressed and functional at the cell surface.

The L42^ICL1^A/S, M43^ICL1^A, and T100^3.44^A mutations served as negative controls (Table S2) and generally did not statistically affect salicin potency (Fig. 3 and Table S3). Only mutation of position 43 to a serine induced a weak decrease of salicin-dependent response in TAS2R16 compared to WT.

The TASR216 I90A/S^3.35^, L91A/S^3.36^, and L185H^5.47^ mutants showed a reduced response to salicin, consistent with their orientation toward the interior of the receptor bundle (Fig. 3 and Table S3). Positions 3.35 and 5.47 have been previously reported to directly interact with ligands^26,30,60^.

Position 239^6.47^ is conserved as Y (64%) and F (8%) in human TAS2Rs (Fig. 4a). In mammals, an aromatic residue (F, Y or H) is also found in 85% of the sequences. Conservation of an aromatic residue also occurs in ORs^16^. The Y239F^6.47^ mutation decreased the potency of salicin by a factor of 11, confirming its importance in receptor activation (Fig. 3). Position Y239^6.47^ corresponded to Y239 and Y241 in TAS2R10 and TAS2R46, respectively. For both of these receptors, the tyrosine to phenylalanine mutation led to a significant reduction in ligand responsiveness. Further, we found that the introduction of an alanine at this position eliminated any response to salicin (data not shown). Born et al. also observed a complete loss of response to agonists with the Y239A^6.47^ TAS2R10 construction^61^. Altogether, these observations highlight the functionnal equivalence of the Y^6.47^FLx motif in TAS2Rs with the F^6.47^YGx in ORs^16^ and the C^6.47^WLP^14^ in non-olfactory class A GPCRs^9^. This motif is particularly important as it forms part of the cradle of the binding pocket and senses the presence of agonists^56^. Adjacent to Y239^6.47^, the aromatic residue F240^6.48^ is conserved as aromatic in 72% of human TAS2Rs and in 67% of mammalian TAS2Rs. As the toggle-switch residue, its nature and function in agonist sensing is similar in ORs (conserved as F^6.48^)^16^ and non-olfactory GPCRs (conserved as W^6.48^)^14^. F240^6.48^ has previously been reported to affect TAS2R16 agonist response. Sakurai et al. showed that mutation of F240^6.48^ to a leucine residue in TAS2R16 drastically alters the function of the receptor, while mutation to aromatic residues (Y and W) leads to moderate changes in the EC50^19^. Further, the potencies of various other agonists were affected in the same manner, highlighting the critical role this residue plays in signal initiation, as is the case for numerous class A GPCRs^14–16^.

**Figure 4.**
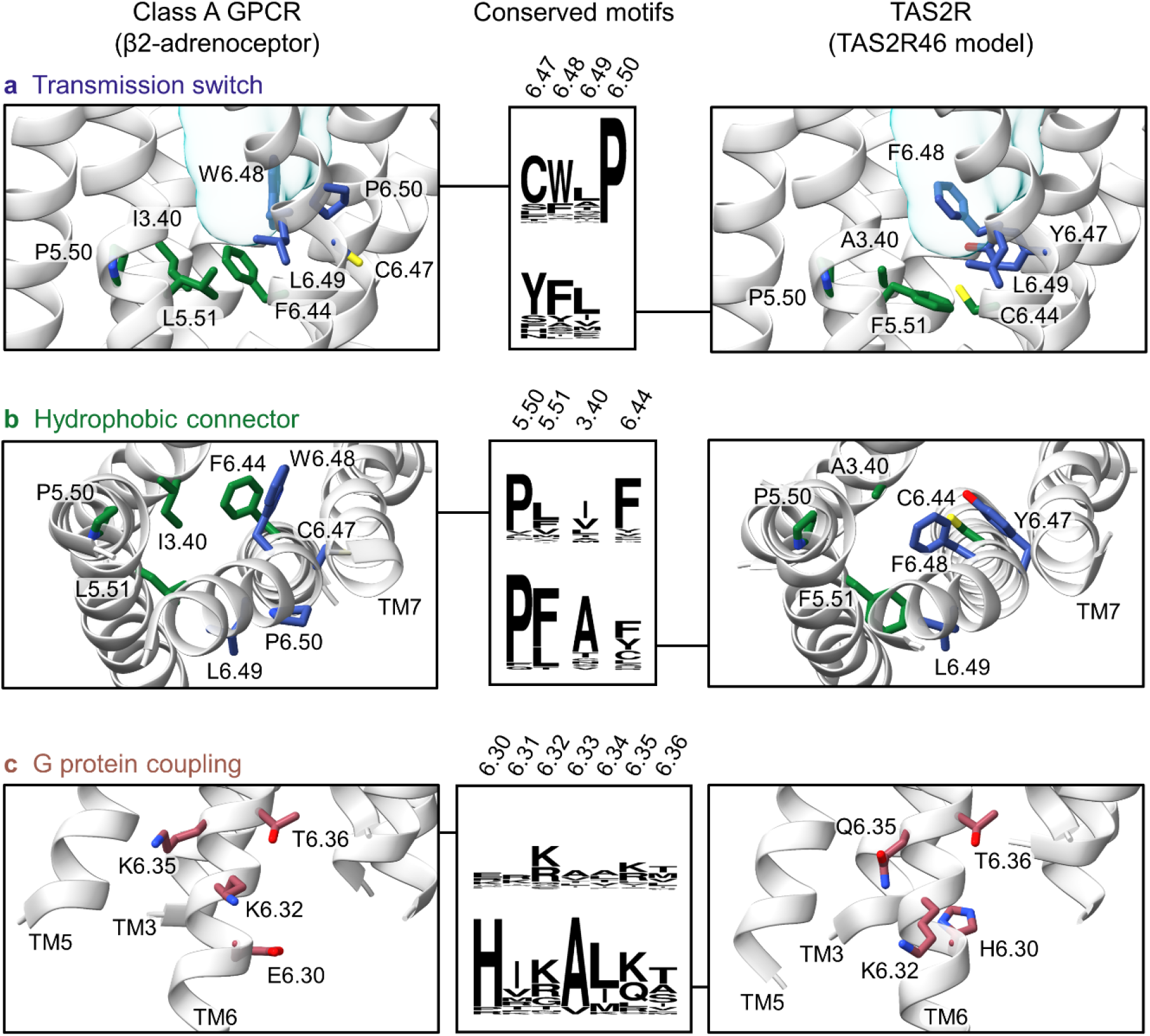
Sequence logos and molecular details of conserved motifs involved in the activation mechanism of class A GPCRs and TAS2Rs, i.e. a) the transmission switch (colored in blue), b) the hydrophobic connector (in green), and c) the G protein-coupling region (in red). The binding pocket is depicted as a pale blue surface. The structure of the β2-adrenoceptor is taken from PDB code 5JQH.

The hydrophobic connector molecular switch involved in class A GPCRs activation^15^ was conserved as P^5.50^I^3.40^F^6.44^ ^14,15,17^. Similarly to other TAS2Rs, a P^5.50^A^3.40^F^6.44^ motif (Fig. 4b) was located at the core of TAS2R16, close to the cradle of the binding pocket. In class A GPCRs, this motif, together with NPxxY^TM7^, holds a central role in receptor signaling, ligand-independent constitutive activation, and β-arrestin signaling in the β2-adrenoceptor^17^. It is plausible that this motif has similar functions in TAS2Rs^62^, as suggested by the modulated response to salicin we found in our mutants (Fig. 3). F236^6.44^, conserved in 75% of mammalian TAS2Rs as Y/F (Fig. 4b), is predicted to be part of the hydrophobic connector molecular switch. The F236A^6.44^ TAS2R16 mutant consistently showed a significantly weaker response to salicin, while no difference in response was found for the F236Q^6.44^ mutant. In a previous study, Thomas et al. found that a F236Y^6.44^ mutation prevented agonist-dependent signaling^25^. In TAS2R14, an alanine residue occupies position 6.44, and mutation to a leucine leads to a decrease in receptor sensitivity to numerous ligands^55^.

Adjacent to position 3.40, S97^3.41^ does not belong to the binding pocket and points toward the membrane. In accordance with a previous report showing its importance for TAS2R16 trafficking^26^, the S97A^3.41^ mutation altered receptor response (gain of function).

Our model predicted that V45^2.39^ is part of a hydrophobic cluster in the intracellular part of TM2 and is conserved as a hydrophobic residue in 72% of TAS2Rs. This hydrophobic area occurs near the highly conserved L229^7.53^ (96% and 93% in humans and mammals, respectively) and the HSFIL^TM7^ motifs and likely forms part of the hydrophobic barrier that prevents flooding of the intracellular region. Mutating V45^2.39^ into a hydrophilic residue (S) strongly altered salicin activation both in this work and in the literature^26^; substitution with a bulkier hydrophobic residue (F) was better tolerated.

In TM6, position 6.29 and adjacent residues have been documented to control G protein selectivity in class A GPCRs^63^. A221^6.29^ and H222^6.30^ are conserved in 60% and 92% of human TAS2Rs, respectively, and in 70% and 94% of mammalian TAS2Rs (Fig. 4c). Position 222^6.30^ is an arginine in TAS2R16. Salicin induced reduced responses in the A221L^6.29^ and R222A^6.30^ mutants, whereas the response of the R222H^6.30^ mutant was not statistically different from the WT. In TASR2R4, the H233A^6.30^ mutation inhibited the response to quinine^64^. Altogether, these findings highlight the need for a positive charge at position 6.30 for G protein-coupling and selectivity.

## Conclusions

This study elucidates key residues and consensus functional motifs of bitter taste receptors (TAS2Rs) using a combination of bioinformatics, molecular modeling, and *in vitro* assays. The consensus sequence motifs match well-known ones in class A GPCRs. Further, we performed sequence alignment of human TAS2Rs with olfactory and non-olfactory class A GPCRs, including residue conservation and experimental data as constraints. Using site-directed mutagenesis, we then evaluated the functional roles of these motifs in TAS2R16 as a case study. In addition to the residues lining the binding pocket, we identified the “toggle/transmission switch” (the YF^6.48^L motif in TM6) and the “hydrophobic connector” (P^5.50^A^3.40^F^6.44^) for agonist sensing. Other molecular switches were identified in the intracellular regions of TM6 and TM7 that are suggested to be involved in G protein selectivity or in receptor activation. These molecular switches extends to mammalian TAS2Rs (see supporting files). The approach, templates, and 3D model provided in this study serve as a foundation for rational design of specific TAS2Rs agonists and antagonists and for decoding sequence-structure-function relationships in these receptors.

## Code and data availability

The scripts used to generate and analyze the models as well as PDB files of TAS2Rs 3D models with the highest meta-score have been deposited on GitHub. (https://github.com/chemosim-lab/TAS2R_data)

## Author contributions

JT^†^, CB^†^, and JP performed numerical modeling. YK and MR conducted functional assays. JT, CB, and SF performed data curation. JT, CB, JP, YK, and MR conducted formal analyses. JT, SF, and JG supervised and managed the study and wrote the paper. MR and JG provided resources for this study.

## Conflicts of interest

The authors declare no competing financial interest.

## Acknowledgements

The authors thank Dr. Xiaojing Cong for fruitful discussion and critical review of the manuscript. This work was funded by the French Ministry of Higher Education and Research [PhD Fellowship to CB], by the National Research Foundation of Korea (NRF) [grant number NRF2020R1A2C2004661], by GIRACT (Geneva, Switzerland) [9th European PhD in Flavor Research Bursaries for first year students to CB], and the Gen Foundation (Registered UK Charity No. 1071026), a charitable trust that primarily funds research in natural sciences, particularly food sciences/technology [grant to CB and JT]. We also benefited from funding by the French government through the UCAJEDI “Investments in the Future” project, managed by the ANR [grant No. ANR-15-IDEX-01 to SF and JG]. Computation for the work described in this paper was supported by the Université Côte d’Azur’s Center for High-Performance Computing.

## Notes

https://github.com/chemosim-lab/TAS2R_data

### Competing Interest Statement

The authors have declared no competing interest.

