## Supplementary Information for "Functional Molecular Switches of Mammalian G Protein-Coupled Bitter-Taste Receptors"

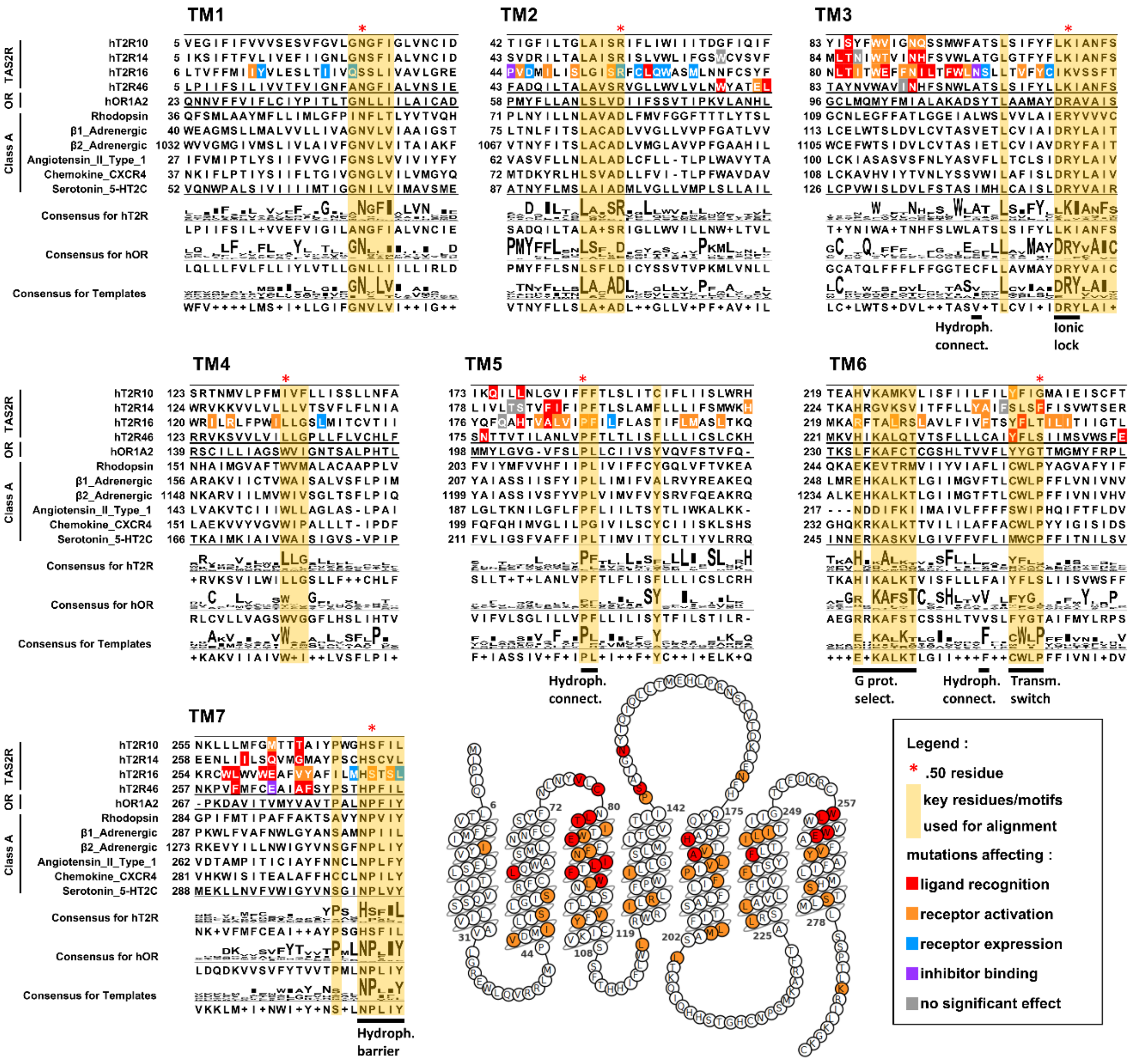

### Figure S1. Alignment and results from site directed mutagenesis.

Selected human type 2 taste receptors (TAS2R), human Olfactory Receptors (OR) and non-OR class A G protein-coupled receptors (GPCRs). In the sequence alignment, residues are colored by their roles as reported in the literature (see legend). The aligned motifs are highlighted in yellow. Consensus sequences for TAS2Rs, ORs and Templates contained 25 human TAS2Rs, 339 human class II ORs, and 6 class A GPCRs, respectively. Functional molecular switches (transmission switch, hydrophobic connector, ionic lock, and hydrophobic barrier) and residues involved in G-protein coupling are indicated under the alignment.

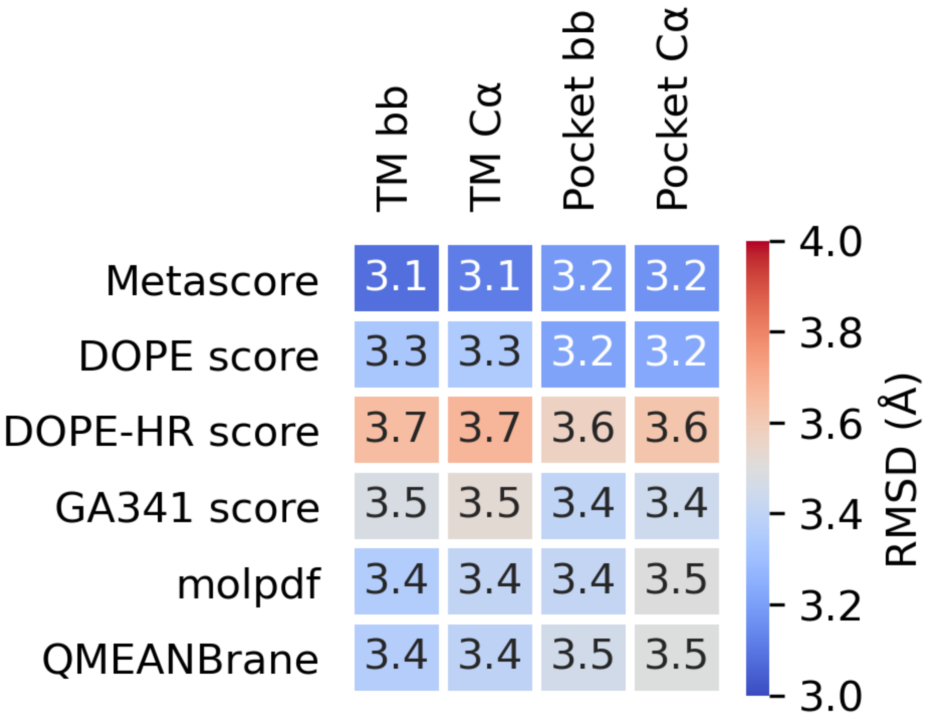

#### Figure S2. RMSD of class F models built using a class A template

The human smoothened receptor (class F) models were built by homology modeling with a class A template (β2-adrenoceptor, PDB 5JQH[1]). The sequence alignment was taken from the GPCRdb[2] and manually refined with UCSF Chimera's structure-based sequence alignment tool (v1.14)[3] based on the 5JQH template and a structure of the smoothened receptor (PDB 4JKV[4]). The same Modeller[5] protocol detailed in the manuscript was used to generate 1000 models of the smoothened receptor. The models were structurally aligned to the 4JKV reference based on the trans membrane (TM) domains and ranked by their meta-scores. Finally, the RMSDs between the reference and each best model were calculated based on the TM domain backbone (TM bb), the TM alpha carbons (TM Ca), the pocket residue backbone (Pocket bb), and the pocket residue alpha carbons (Pocket Ca). The pocket residues were identified by visual inspection of four class F X-ray structures in complex with a ligand (PDB codes 6O3C[6], 4JKV[4], 4QIM[7], and 4N4W[7]).

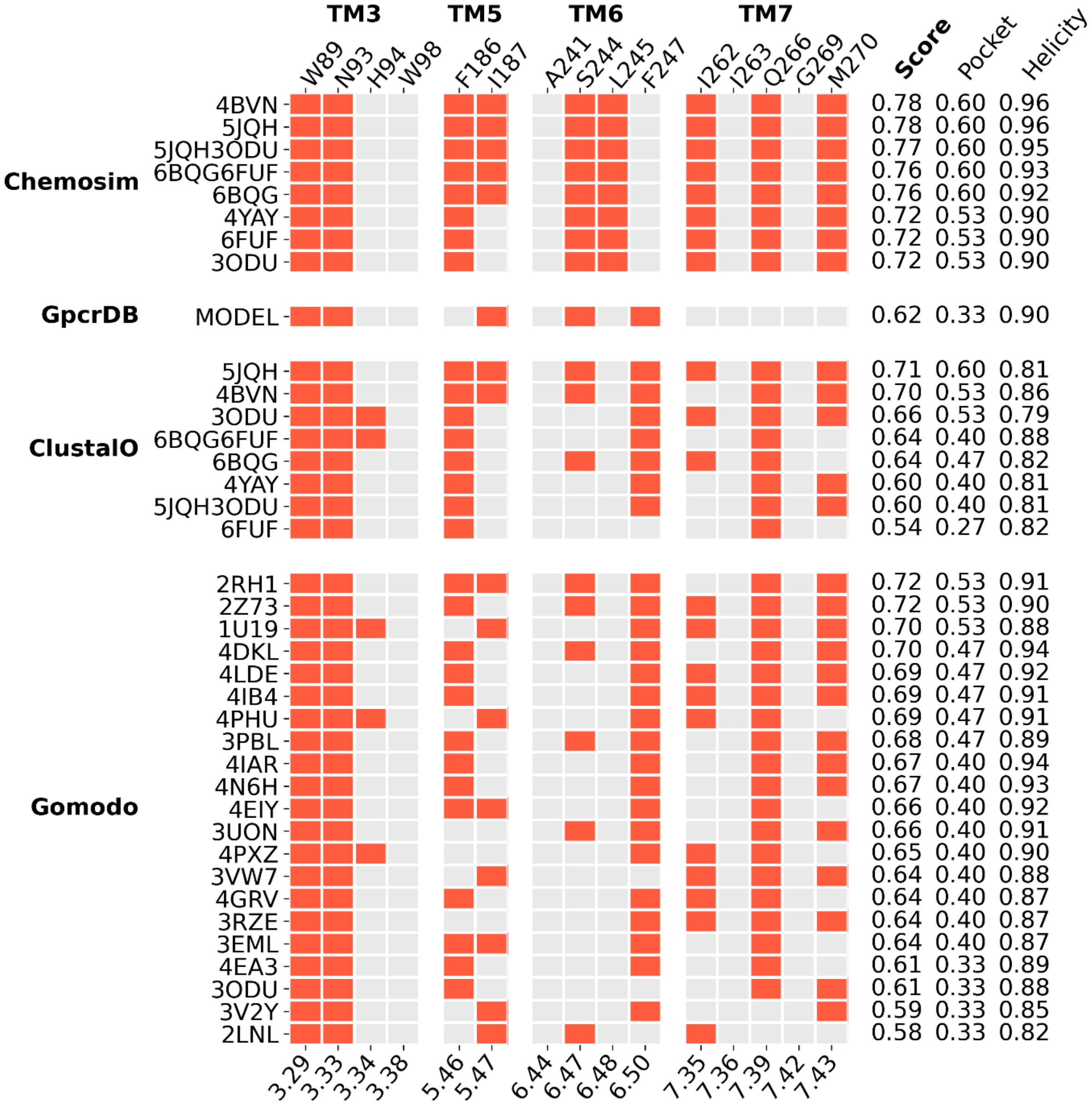

#### Figure S3.a. Detailed analysis of TAS2R14 binding pocket residues

Meta-scores of top models for each protocol and template. Best models following the Gomodo[8] and ClustalO[9] protocols were selected based on their DOPE score[10]. For BitterDB[11], the only available model did not satisfy our structure quality criteria. The x-axis labels correspond to the Ballesteros-Weinstein numbering of each residue[12]. The left y-axis provides the PDB code of each template except for GPCRdb, where the model was retrieved directly from their website. The right y-axis shows the meta-score, pocket score, and helicity score for each selected model.

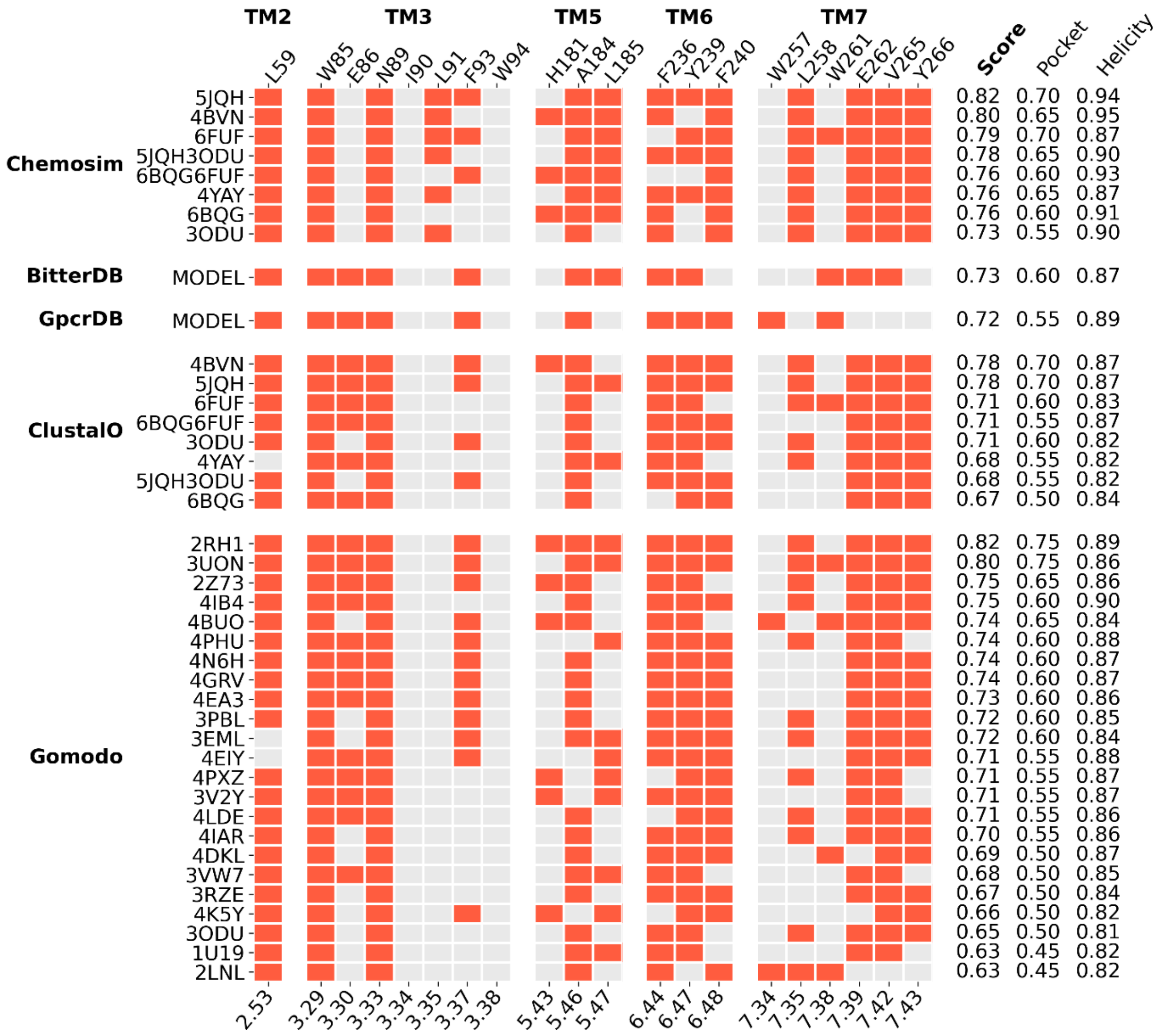

### Figure S3.b. Detailed analysis of TAS2R16 binding pocket residues

Meta-scores of top models for each protocol and template. Best models following the Gomodo and ClustalO protocols were selected based on their DOPE score. The x-axis labels correspond to the Ballesteros-Weinstein numbering of each residue. The left y-axis provides the PDB code of each template except for BitterDB and GPCRdb, where the model was retrieved directly from their website. The right y-axis shows the meta-score, pocket score, and helicity score for each selected model.

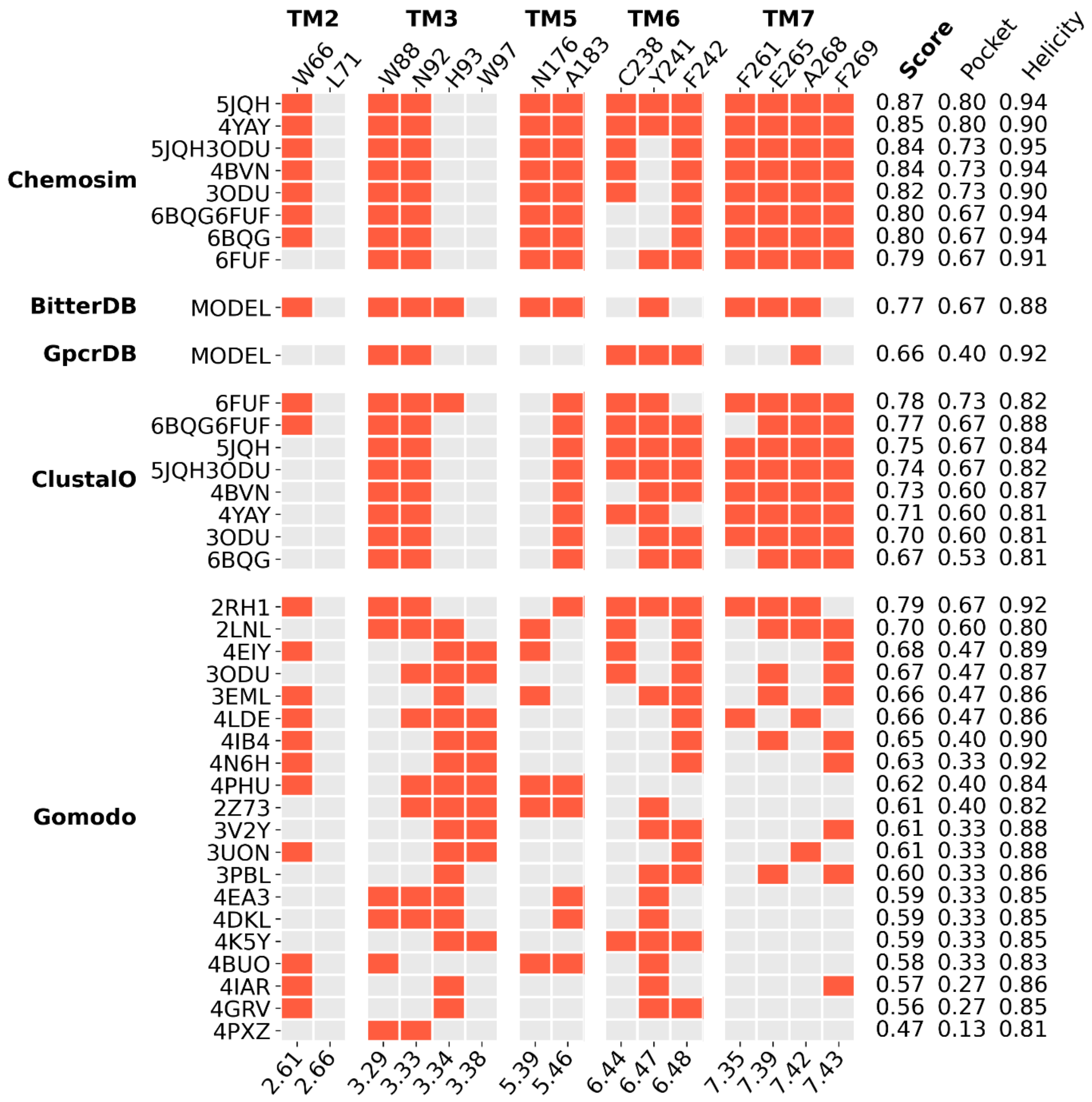

#### Figure S3.c. Detailed analysis of TAS2R46 binding pocket residues

Meta-scores of top models for each protocol and template. Best models following the Gomodo and ClustalO protocols were selected based on their DOPE score. The x-axis labels correspond to the Ballesteros-Weinstein numbering of each residue. The left y-axis provides the PDB code of each template except for BitterDB and GPCRdb, where the model was retrieved directly from their website. The right y-axis shows the meta-score, pocket score, and helicity score for each selected model.

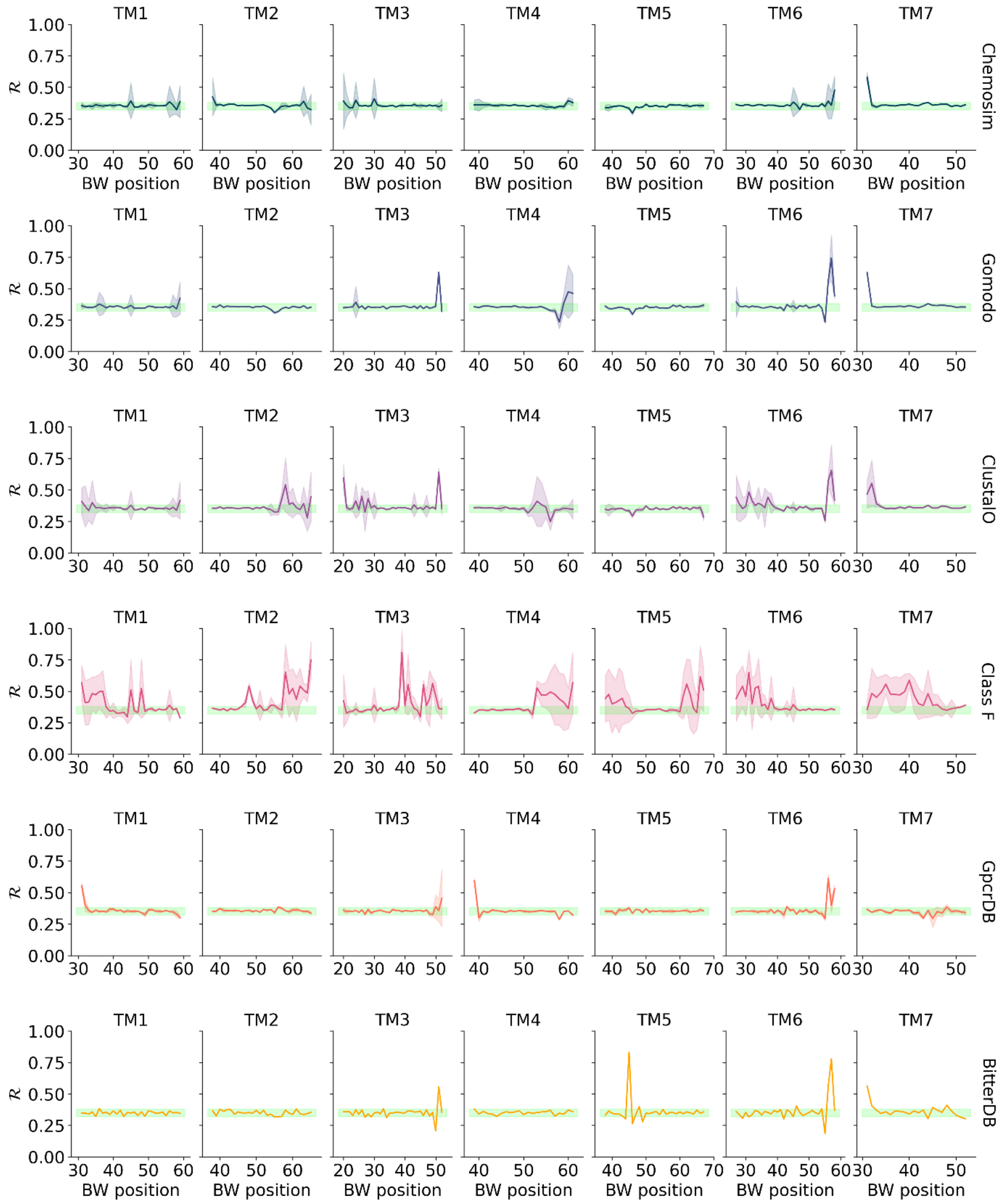

### Figure S4.a. Analysis of TAS2R14 transmembrane helicity

Ramachandran number *(R)* plot of each residue, numbered by their Ballesteros-Weinstein (BW) position, for the models produced by the best template for each protocol. Standard deviation is represented by the shaded area, and the green zone corresponds to *R* values typically found in alpha helices of crystallographic GPCR structures (0.32 to 0.38).

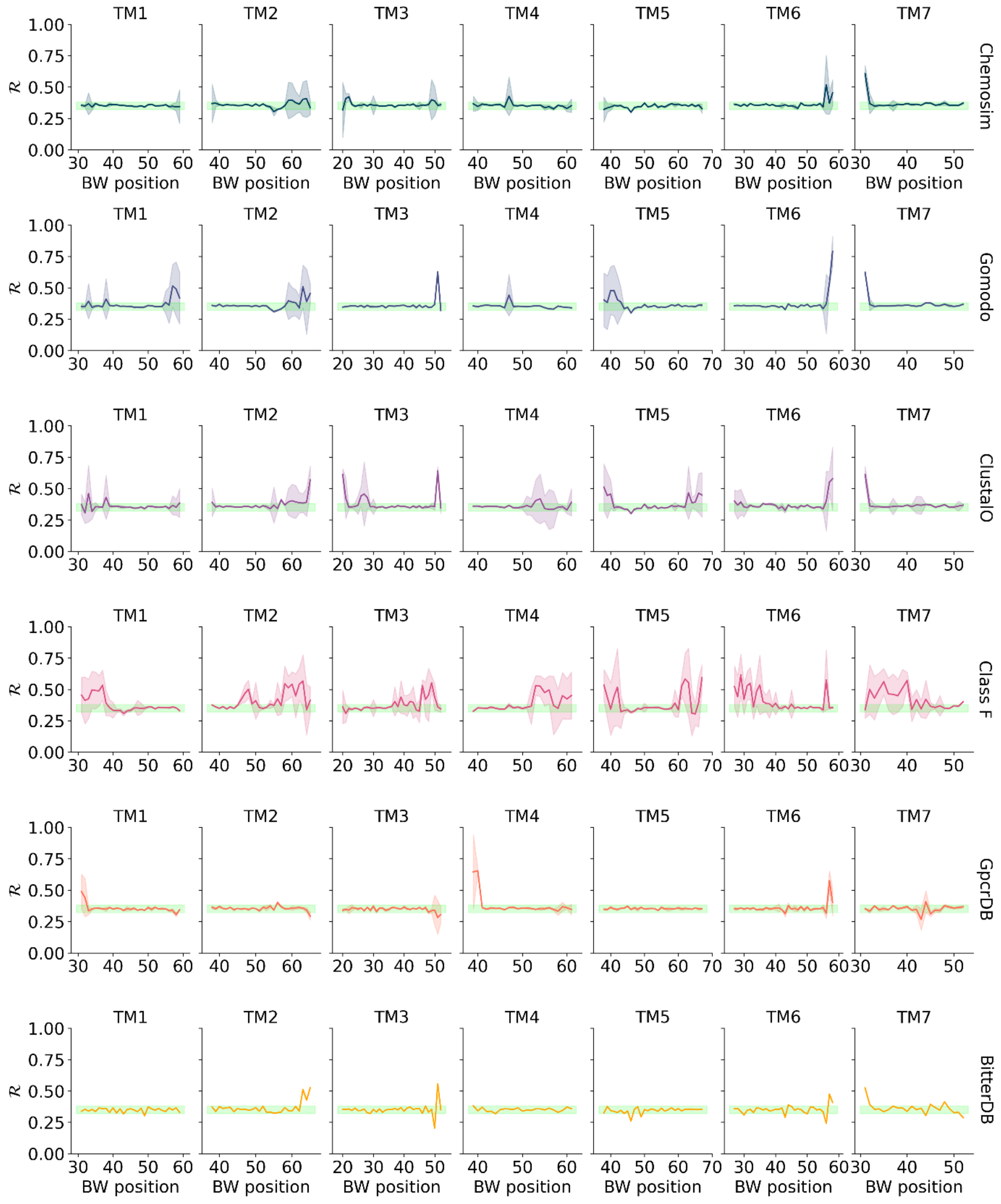

### Figure S4.b. Analysis of TAS2R16 transmembrane helicity

See figure caption S4.a.

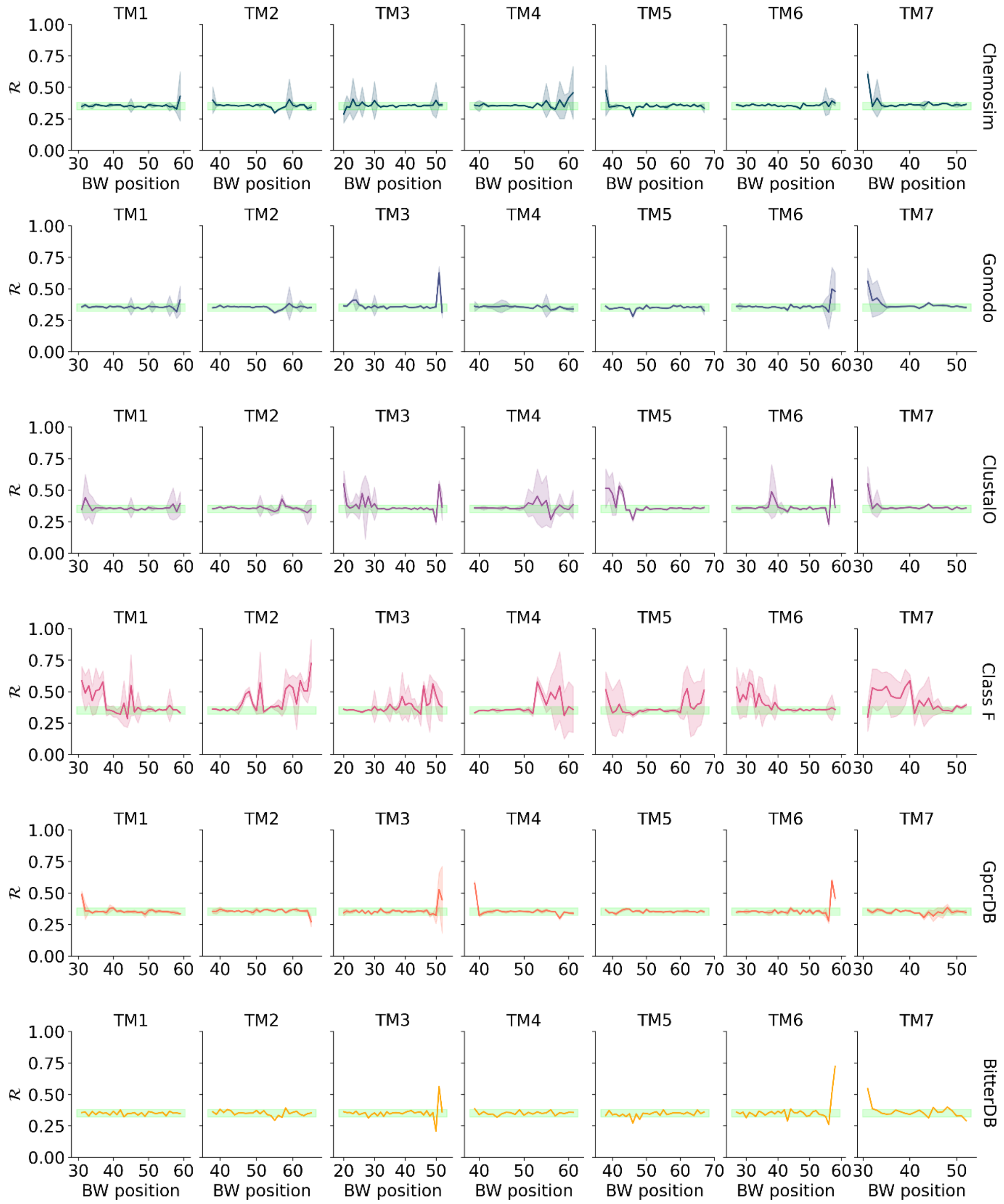

### Figure S4.c. Analysis of TAS2R46 transmembrane helicity

See figure caption S4.a.

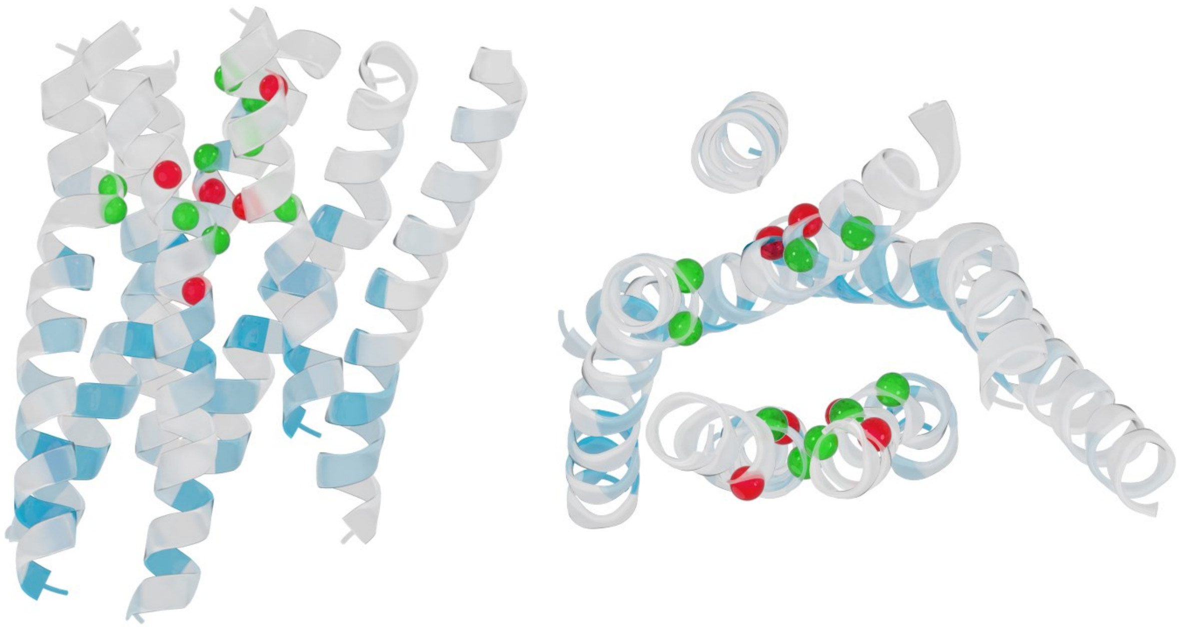

#### Figure S5.a. Structure of the TAS2R14 model with the highest meta-score

Structure of the best *Chemosim* model obtained from the present study. The residues defining the binding pocket are shown as spheres if their side chains are oriented outward (red) or inward (green) from the pocket and follow from the results shown in Fig S3. Positions of the highly conserved residues in the human TAS2R family are indicated by a color scale, from 50% or less conservation (white) to 100% (blue).

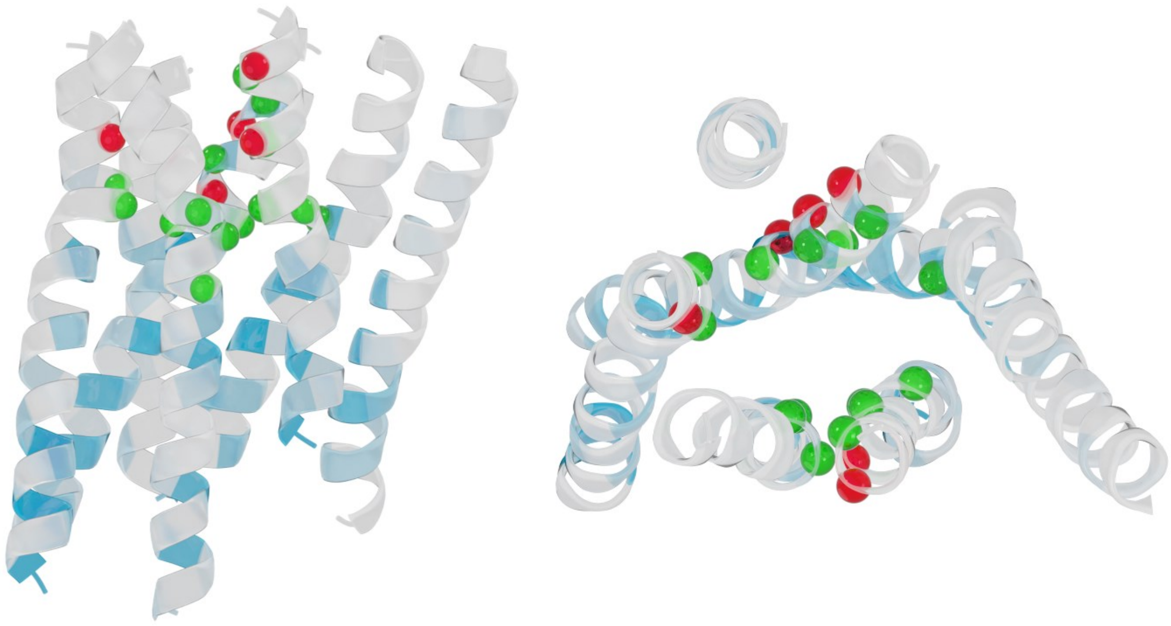

#### Figure S5.b. Structure of the TAS2R16 model with the highest meta score

See figure caption S5.a.

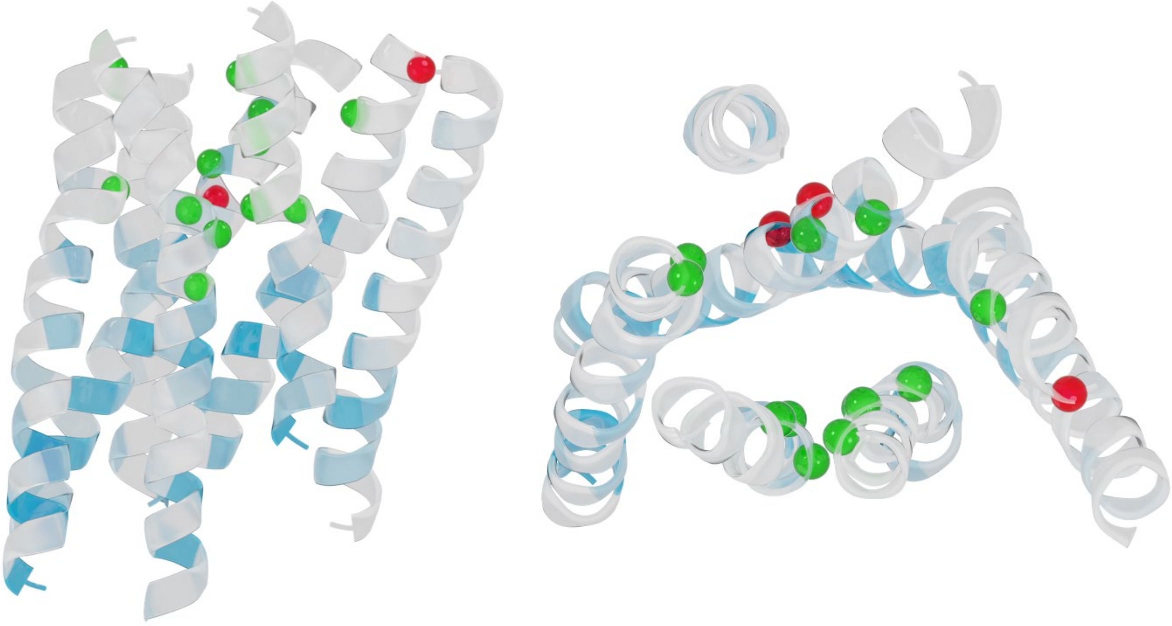

#### Figure S5.c. Structure of the TAS2R46 model with the highest meta score

See figure caption S5.a.

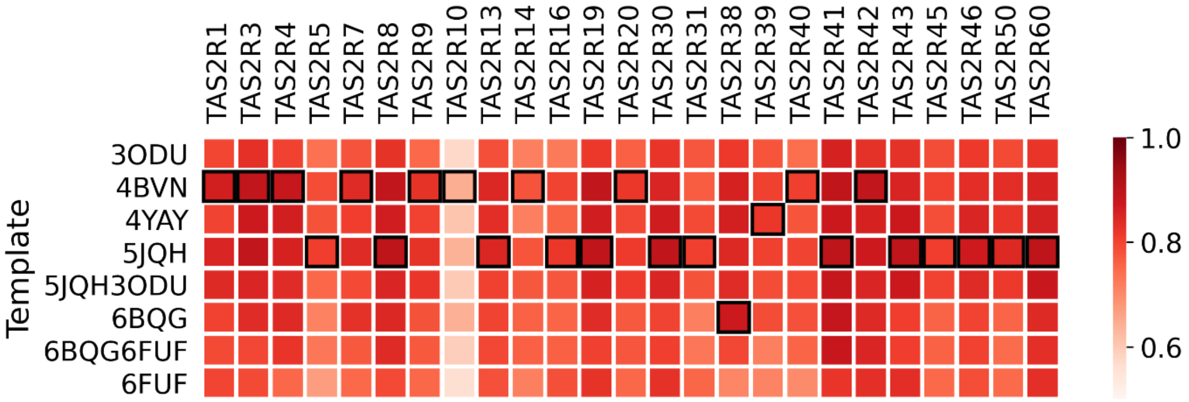

#### Figure S6. Selection of TAS2R models according to various class A templates

TAS2R models were built following the *Chemosim* protocol. The best models are shown with black boxes and were selected according to the highest meta-score. For all receptors, a consensus TAS2R cavity was used for the detection of residues oriented in the binding pocket. This consensus cavity was composed of residues 3.29, 3.33, 3.34, 3.38, 5.46, 6.44, 6.47, 6.48, 7.35, 7.39, 7.42, and 7.43 and was completed by receptor-specific cavity residues highlighted in the annotated TAS2Rs MSA that is provided in the supplementary files (TAS2R-msa-annotated.xlsx).

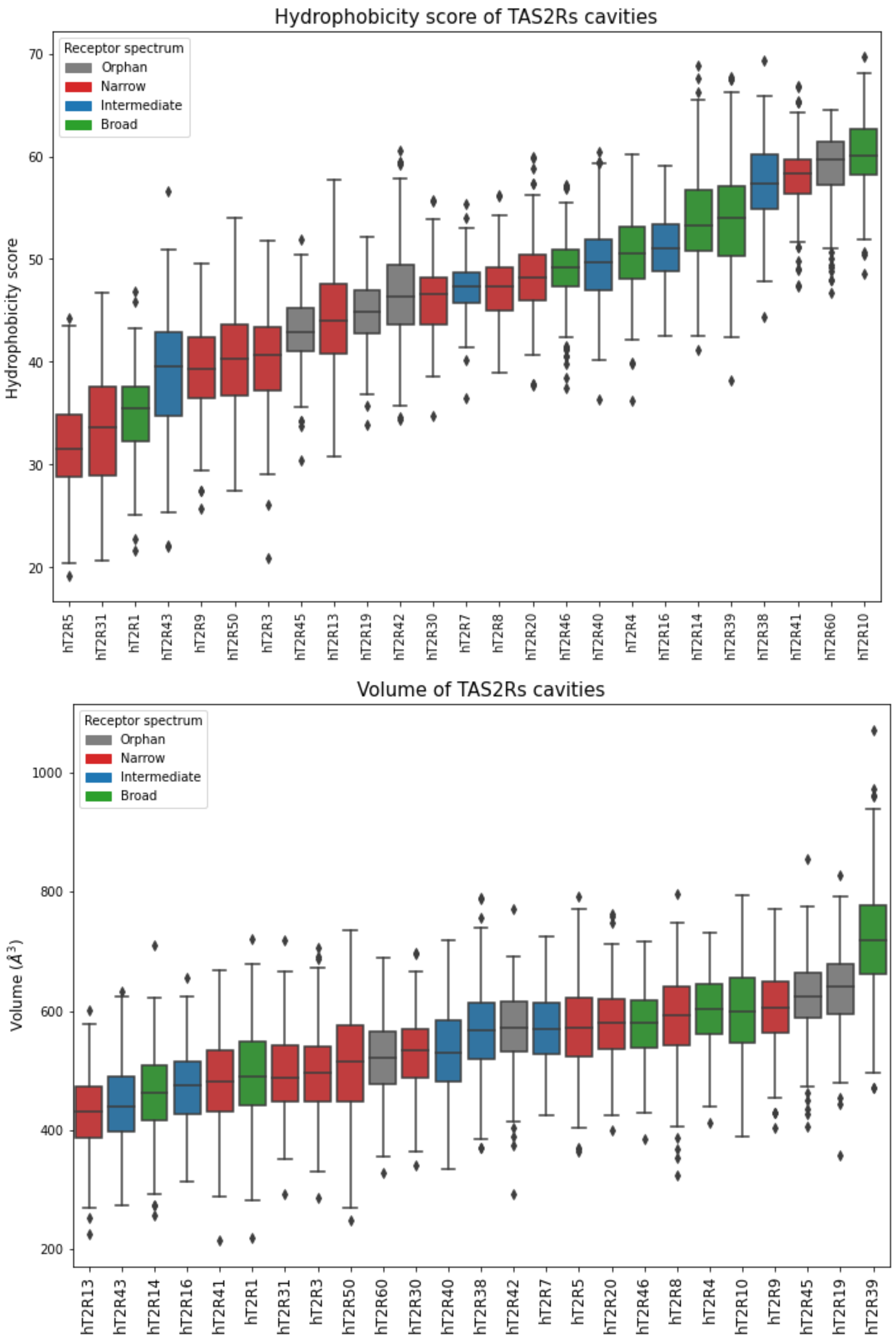

#### Figure S7. Structural analysis of TAS2R binding pocket

Box-plot of hydrophobicity and volume of TAS2Rs binding pocket. The box extends from the lower to upper quartile values of the data, with a line at the median and outliers plotted in diamonds. The top 250 models for each TAS2R produced by the *Chemosim* protocol and selected templates as shown in Figure S6 were analyzed by MDpocket[13] and colored according to the receptive range (broad, intermediate/specific, narrow, and orphan receptors in green, blue, red, and grey, respectively). A positive hydrophobicity score means that the cavity is mainly hydrophobic.

### Table S1. Summary of the most conserved TAS2R amino acids

The most conserved TAS2R residues (above 80% sequence identity) and those involved in TAS2R hallmarks (in yellow/bold) used for multiple sequence alignment with OR and class A templates.

| | ClassA motif | OR motif | TAS2R Motif | TAS2R Consensus | Conservation | BW numbering | TAS2R14 | TAS2R16 | TAS2R46 |
|---|---|---|---|---|---|---|---|---|---|
| **TM1** | | | | G | 88% | 1.46 | G20 | I21 | G20 |
| | **GNxLV** | **GNLLI** | **NGFI** | **N** | **92%** | **1.50** | N24 | S25 | N24 |
| | | | | **G** | **72%** | **1.51** | S25 | S26 | G25 |
| | | | | **F** | **84%** | **1.52** | F26 | L27 | F26 |
| | | | | **I** | **92%** | **1.53** | I27 | I28 | I27 |
| **ICL1** | | | | W | 80% | | W35 | W36 | W35 |
| **TM2** | | | | D | 84% | 2.40 | D45 | D46 | D45 |
| | | | | I | 84% | 2.42 | I47 | I48 | I47 |
| | | | | L | 80% | 2.43 | L48 | L49 | L48 |
| | **LAxAD** | **LSxxD** | **LAxSR** | **L** | **100%** | **2.46** | L51 | L52 | L51 |
| | | | | **A** | **64%** | **2.47** | A52 | G53 | A52 |
| | | | | **S** | **84%** | **2.48** | S54 | S55 | S54 |
| | | | | **R** | **96%** | **2.49** | R55 | R56 | R55 |
| | | | | L | 92% | 2.53 | L58 | L59 | L58 |
| **TM3** | | | | W | 84% | 3.29 | W89 | W85 | W88 |
| | | | | N | 84% | 3.33 | N93 | N89 | N92 |
| | | | | W | 100% | 3.38 | W98 | W94 | W97 |
| | **L** | **L** | **L** | **L** | **96%** | **3.43** | L103 | L99 | L102 |
| | | | | F | 80% | 3.46 | F106 | F102 | F105 |
| | | | | Y | 92% | 3.47 | Y107 | Y103 | Y106 |
| | **DRY** | **MAYDRYVAIC** | **KIANFS** | **K** | **92%** | **3.50** | K110 | K106 | K109 |
| | | | | **I** | **88%** | **3.51** | I111 | V107 | I110 |
| | | | | **A** | **76%** | **3.52** | A112 | S108 | A111 |
| | | | | **N** | **64%** | **3.53** | N113 | S109 | N112 |
| | | | | **F** | **84%** | **3.54** | F114 | F110 | F113 |
| | | | | **S** | **64%** | **3.55** | S115 | T111 | S114 |
| **ICL2** | | | | F | 88% | | F119 | F115 | F118 |
| | | | | L | 88% | | L122 | L118 | L121 |
| **TM4** | | | | K | 84% | 4.39 | K123 | R119 | K122 |
| | **W** | **W** | **LLG** | **L** | **88%** | **4.50** | L134 | L130 | L133 |
| | | | | **L** | **80%** | **4.51** | L135 | L131 | L134 |
| | | | | **G** | **72%** | **4.52** | V136 | G132 | G135 |
| **ECL2** | | | | N | 100% | | N162 | N163 | N161 |
| | | | | T | 96% | | T164 | T165 | T163 |
| **TM5** | **P** | **PF** | **PF** | **P** | **92%** | **5.50** | P190 | P188 | P187 |
| | | | | **F** | **72%** | **5.51** | F191 | F189 | F188 |
| | | | | L | 80% | 5.55 | L195 | L193 | L192 |
| | **Y** | **Y** | **F** | **F** | **52%** | **5.58** | F198 | T196 | F195 |
| | | | | L | 100% | 5.61 | L201 | L199 | L198 |
| | | | | S | 100% | 5.64 | L204 | S202 | L201 |
| | | | | L | 96% | 5.65 | M205 | L203 | L202 |
| **ICL3** | | | | H | 96% | | H208 | Q206 | H205 |
| | | | | G | 84% | | I218 | G213 | I215 |
| | | | | D | 84% | | D221 | N216 | D218 |
| | | | | P | 80% | | A222 | P217 | P219 |
| **TM6** | **KxxK** | **RxKAFSTC** | **HxKALKT** | **H** | **92%** | **6.30** | H227 | R222 | H224 |
| | | | | **K/R** | **60%** | **6.32** | G229 | T224 | K226 |
| | | | | **A** | **92%** | **6.33** | V230 | A225 | A227 |
| | | | | **L** | **64%** | **6.34** | K231 | L226 | L228 |
| | | | | **K/Q** | **88%** | **6.35** | S232 | R227 | Q229 |
| | | | | **T/S** | **60%** | **6.36** | V233 | S228 | T230 |
| | | | | F | 96% | 6.40 | F237 | L232 | F234 |
| | | | | L | 80% | 6.43 | Y240 | V235 | L237 |
| | **CWLP** | **FYG** | **YFL** | **Y** | **64%** | **6.47** | S244 | Y239 | Y241 |
| | | | | **F** | **60%** | **6.48** | L245 | F240 | F242 |
| | | | | **L/I/V** | **76%** | **6.49** | S246 | L241 | L243 |
| **TM7** | **NPxxY** | **PxxNPxIY** | **PxxHSFIL** | **P** | **76%** | **7.46** | P273 | I269 | P272 |
| | | | | **H** | **96%** | **7.49** | H276 | H272 | H275 |
| | | | | **S** | **68%** | **7.50** | S277 | S273 | P276 |
| | | | | **F** | **60%** | **7.51** | C278 | T274 | F277 |
| | | | | **I** | **76%** | **7.52** | V279 | S275 | I278 |
| | | | | **L** | **96%** | **7.53** | L280 | L276 | L279 |
| | | | | I | 92% | 7.54 | I281 | M277 | I280 |
| | | | | N | 80% | | N284 | S280 | N283 |
| | | | | L | 96% | | L287 | L283 | L286 |

#### Table S2. Mutations tested *in vitro* to assess the 3D model

| Mutations | TAS2R motifs | Location/role |
|---|---|---|
| I90A/S$^{3.35}$, L91A/S$^{3.36}$, L185A$^{5.47}$ | n.a. | inside pocket |
| T100A$^{3.44}$ | Negative control | outside pocket |
| S97A/N$^{3.41}$ | n.a. | receptor surface/ receptor trafficking |
| F236A/Q$^{6.44}$ | P$^{5.50}$A$^{3.40}$F$^{6.44}$ | pocket cradle/ hydrophobic connector, agonist sensing |
| Y239F$^{6.47}$ | YF$^{6.48}$L | pocket cradle/ transmission switch, agonist sensing |
| V45S/F$^{2.39}$ | Next to PxxHS$^{7.50}$FIL | intracellular part/ hydrophobic barrier |
| L42A/S$^{ICL1}$, M43A/S$^{ICL1}$ | Negative control | intracellular part |
| A221L$^{6.29}$, R222A/H$^{6.30}$ | HxK$^{6.32}$ALKT | G protein binding site/ G protein selectivity |

#### Table S3. Salicin-induced *in vitro* response in wild-type and mutant TAS2R16

| | Mutations | $EC_{50}$ † (mM) | Maximal Response ($\Delta F/F_0$) |
|---|---|---|---|
| WT | | 0.98 ± 0.01 | 0.55 |
| I90 | I90A | 3.34 ± 0.03*** | 0.50 |
| | I90S | 3.20 ± 0.11*** | 0.31 |
| L91 | L91A | 2.85 ± 0.03*** | 0.57 |
| | L91S | 6.05 ± 0.03*** | 0.47 |
| L42 | L42A | 0.61 ± 0.04 | 0.37 |
| | L42S | 1.23 ± 0.05 | 0.33 |
| M43 | M43A | 0.53 ± 0.12 | 0.45 |
| | M43S | 1.77 ± 0.13** | 0.40 |
| V45 | V45S | 3.30 ± 0.12*** | 0.41 |
| | V45F | 2.79 ± 0.12** | 0.26 |
| S97 | S97A | 0.17 ± 0.04*** | 0.50 |
| | S97N | 0.92 ± 0.11 | 0.31 |
| T100 | T100A | 0.50 ± 0.06 | 0.61 |
| L185 | L185H | 3.87 ± 0.05*** | 0.27 |
| A221 | A221L | 3.78 ± 0.04*** | 0.38 |
| R222 | R222A | 5.10 ± 0.08*** | 0.34 |
| | R222H | 0.69 ± 0.10 | 0.52 |
| F236 | F236A | 10.38 ± 0.11*** | 0.39 |
| | F236Q | 0.57 ± 0.08 | 0.50 |
| Y239 | Y239F | 11.30 ± 0.08*** | 0.42 |

† Values are means ± SEM; Statistical significance is indicated by *** $P < 0.001$, ** $P < 0.01$, and * < $P$ 0.05 vs. the WT group (one-way ANOVA followed by Dunnett's test)

#### Other supporting information files

The MSA of human TAS2Rs and a selection of ORs and class A templates (**TAS2R-OR-templates.pir**); the MSA of reviewed mammalian TAS2R sequences obtained from Uniprot (**mammalian-TAS2R.pir**); and an annotated MSA of human TAS2Rs (**TAS2R-msa-annotated.xlsx**).
